# Preclinical Prediction of Resistance and Optimization of Sequential Therapy for ALK-positive Lung Cancer Using Next-generation ALK Inhibitors

**DOI:** 10.1101/2025.05.26.656085

**Authors:** Yuki Takei, Hirotaka Kuroiwa, Chisaki Arai, Yuta Doi, Kentaro Semba

**Affiliations:** Department of Life Science and Medical Bioscience, Graduate School of Advanced Science and Engineering, Waseda University, 2-2 Wakamatsu-cho, Shinjuku-ku, Tokyo 162-8480, Japan; Translational Research Center, Fukushima Medical University, Hikarigaoka, Fukushima 960-1295, Japan

## Abstract

**Background:** Anaplastic lymphoma kinase (ALK) gene rearrangements occur in approximately 5% of non-small cell lung cancers (NSCLCs). Although ALK tyrosine kinase inhibitors provide substantial clinical benefits, acquired resistance-conferring mutations frequently emerge, leading to disease progression. Preclinical prediction of these mutations might help guide the development of more effective sequential treatment strategies prior to clinical application.

**Objective:** To predict the emergence of resistance mutations to the investigational ALK inhibitors zotizalkib (TPX-0131), gilteritinib (ASP2215), and neladalkib (NVL-655) following resistance to first-line alectinib and assess the potential of these drugs as second-line therapies.

**Methods:** A polymerase chain reaction (PCR)-based mutagenesis system was used to introduce random mutations into ALK cDNA harboring representative alectinib-resistant mutations. Mutant libraries were expressed in Ba/F3 cells, which were exposed to each inhibitor. Drug-resistant clones were isolated, sequenced, and evaluated for drug sensitivity using viability assays and immunoblotting.

**Results:** Several resistance mutations against zotizalkib, gilteritinib, and neladalkib were identified. Sequential use of these agents effectively suppressed all predicted resistance patterns with G1202R or I1171N.

**Conclusions:** This PCR-based platform provides a valuable approach for anticipating resistance mutations and guiding the design of optimized sequential therapies. Zotizalkib, gilteritinib, and neladalkib might represent promising alternatives to lorlatinib as second-line treatments for ALK-positive NSCLC.

**Key Points:**

- PCR-based mutation prediction system was successfully applied to fourth-generation ALK inhibitors.
- showed efficacy against G1202R-positive relapses with minimal evidence of secondary resistance mutations.
- combinations of gilteritinib with either neladalkib or ensartinib may sustain efficacy and delay resistance in I1171N-positive relapses.

## 1 Introduction

The anaplastic lymphoma kinase (ALK) gene has been recognized as a tumor-agnostic oncogene, and the term “ALKoma” has been proposed to reflect the oncogenic activation of ALK across several malignancies. ALK activation occurs through multiple mechanisms, including point mutations, gene amplification, and chromosomal rearrangement [1]. Lung cancer remains the leading cause of cancer-related mortality globally, accounting for approximately 1.8 million deaths annually, and ALK rearrangements have been identified in approximately 5% of non-small cell lung cancers (NSCLCs) [2, 3]. In most cases of ALK-rearranged NSCLC, an inversion on chromosome 2 leads to the fusion of the ALK gene with the echinoderm microtubule-associated protein-like 4 (EML4) gene [4]. The resulting EML4– ALK fusion protein remains constitutively active through the oligomerization (coiled-coil) domain of EML4, driving downstream signaling pathways that promote uncontrolled cell proliferation and tumor progression [5–7].

Multiple ALK tyrosine kinase inhibitors (ALK-TKIs) have been developed for ALK-positive NSCLC. Crizotinib, the first ALK-TKI approved for clinical use, achieved better progression-free survival than standard chemotherapy in phase III trials [8]. Subsequently, alectinib, a second-generation ALK-TKI, exhibited even greater efficacy, and it has become the preferred first-line treatment for ALK-positive NSCLC [9]. However, resistance commonly develops within a few years, often because of secondary point mutations in the ALK kinase domain such as G1202R or I1171N [10].

To overcome resistance mutations, the third-generation ALK-TKI lorlatinib was developed, and it is currently recommended as a second-line treatment following alectinib failure [11, 12]. Nevertheless, compound mutations, such as G1202R + L1196M, can emerge after lorlatinib treatment, leading to further resistance and disease progression [13, 14]. In addition, lorlatinib has been linked to frequent and severe adverse effects, including hypercholesterolemia, hypertriglyceridemia, peripheral neuropathy, and cognitive impairment. These toxicities are generally more pronounced than those observed with other ALK-TKIs, thereby compromising patients’ quality of life [15]. Furthermore, some patients exhibit primary resistance to lorlatinib, underscoring the need for alternative second-line strategies. In this context, anticipating resistance mutations prior to treatment represents a promising approach to guiding the selection of effective sequential treatments.

Several groups developed resistance prediction systems using N-ethyl-N-nitrosourea (ENU)-based mutagenesis screens [14, 16]. However, ENU induces mutations through chemical modification of DNA bases, leading to a biased mutation spectrum and potentially incomplete coverage of relevant variants [17]. Furthermore, because ENU introduces genome-wide mutations, it is difficult to assess the effects of ALK mutations in isolation.

To address these limitations, we previously established a resistance prediction platform using error-prone polymerase chain reaction (PCR), which specifically introduces mutations into the ALK kinase domain. Using this system, we predicted resistance mutations that arise when repotrectinib or ensartinib is used as a second-line treatment against alectinib-resistant variants such as G1202R-and I1171N-mutated cancer [18]. However, the use of repotrectinib or ensartinib led to the emergence of mutations that could not be treated with existing drugs. Therefore, it is necessary to explore compounds, including fourth-generation ALK inhibitors, capable of preventing the emergence of untreatable resistance mutations.

In this study, we applied an error-prone PCR-based mutagenesis approach to three compounds, including the fourth-generation ALK inhibitors zotizalkib (TPX-0131) and neladalkib (NVL-655), as well as gilteritinib (ASP2215), which was originally developed for acute myeloid leukemia and is currently being evaluated in clinical trials for ALK-positive NSCLC. Our aim was to predict resistance mutations that can emerge when these compounds are used following alectinib failure. Furthermore, we assessed whether other ALK-TKIs could overcome resistance mutations that were predicted to arise after second-line treatment with zotizalkib, gilteritinib, or neladalkib and evaluated their potential utility as alternative second-line therapeutic options to lorlatinib.

## 2 Materials and Methods

### 2.1 Cell Lines and Culture Condition

Ba/F3 cells, which are murine bone marrow-derived pro-B cells, were cultured in RPMI-1640 medium (FUJIFILM Wako, Osaka, Japan) supplemented with 10% fetal bovine serum (FBS; Nichirei Biosciences, Tokyo, Japan), 100 U/mL penicillin (Meiji Seika Pharma, Tokyo, Japan), 100 µg/mL streptomycin (Meiji Seika Pharma), and 10 ng/mL murine interleukin 3 (IL-3; PeproTech, Cranbury, NJ, USA) and incubated at 37℃ in a 5% CO_2_ atmosphere. Platinum-E retroviral packaging cells (Plat-E cells) were cultured in Dulbecco’s Modified Eagle Medium (low glucose; FUJIFILM Wako) supplemented with 10% FBS, 100 U/mL penicillin, and 100 µg/mL streptomycin and incubated at 37℃ and 5% CO_2_.

### 2.2 Reagents

Alectinib (CH5424802), crizotinib (PF-02341066), lorlatinib (PF-06463922), neladalkib (NVL-655), and brigatinib (AP26113) were purchased from Selleck Chemicals (Houston, TX). Zotizalkib (TPX-0131) and gilteritinib (ASP2215) were purchased from MedChemExpress (Monmouth Junction, NJ, USA). All inhibitors were dissolved in dimethyl sulfoxide (DMSO) and stored at −80℃ after aliquoting.

### 2.3 Construction of Complementary DNA (CDNA) Mutant Libraries Using Error-prone PCR

We previously generated cDNA mutant libraries possessing G1202R or I1171N and a random mutation [18]. These constructs were generated by cloning the cDNA sequence of EML4–ALK variant 1 (GenBank: AB274722.1), which is the most common variant, being observed in approximately 45% of ALK-positive NSCLCs [19], into the pMXs-GW-IRES-Puro vector. The inserted sequence harbored either the G1202R or I1171N mutation. Error-prone PCR targeting the ALK kinase domain was subsequently performed to introduce random mutations. We used this library for this study. Detailed methods were described previously [18].

### 2.4 Establishment of Ba/F3 Cells Expressing cDNA Mutant Libraries

For retrovirus production, Plat-E cells (2 × 10^6^ cells) were seeded into a 100-mm cell culture dish. After overnight culture, 10 μg of the established cDNA mutant library and 30 μg of polyethylenimine were added to the culture. After incubation for 8 h, the medium was replaced with RPMI-1640. After another 24 h of incubation, the cell culture containing retrovirus was harvested, and cell debris was removed via centrifugation for 15 min at 3500 rpm. Cultured Ba/F3 cells (1 mL; 5 × 10^6^ cells/well, three wells in total) were seeded into 12-well plates and mixed with 1 mL of the harvested retrovirus supplemented with 8 μg/mL polybrene for spinfection. The Ba/F3 cells were centrifuged in the 12-well plates for 1 h at 32℃ (900 × *g*). Thereafter, all cultures were transferred to 25-cm^2^ cell culture flasks, and 9 mL of the mixture of RPMI-1640 medium and the harvested virus supplemented with 8 μg/mL polybrene were added. After overnight incubation at 37℃ and 5% CO_2_, the virus was removed via washing and centrifugation, and the cells were transferred to 30 mL of RPMI-1640 medium supplemented with 0.5 ng/mL IL-3 in a 75-cm^2^ culture flask. After 24 h of incubation, the culture medium was changed to 60 mL of RPMI-1640 supplemented with 0.05 ng/mL IL-3 and 1 μg/mL puromycin (FUJIFILM Wako) in a 75-cm^2^ culture flask. Puromycin was applied to Ba/F3 cells for 48 h before establishing the cDNA mutant library of Ba/F3 cells. To assess viral infection efficiency, the infected cells were transferred to six-well plates (2 × 10^5^ cells /mL in 2 mL of culture), and puromycin (final concentration 1 μg/mL) was then added to each well individually. Living cells were monitored using a hemocytometer by staining dead cells with 0.5% trypan blue over 40 h after the addition of puromycin when non-infected cells were completely dead. The infection efficiency was calculated by drawing the growth curve of the cells and predicting the percentage of infected cells against the total number of living cells before the addition of puromycin.

### 2.5 Identification of Resistance Mutations Against Zotizalkib, Gilteritinib, or Neladalkib

Ba/F3 cells (1000 cells) expressing EML4–ALK and carrying G1202R and a random mutation were seeded into 96-well plates and cultured with zotizalkib (100 nM) for 2 weeks. Ba/F3 cells (1000 cells) expressing EML4–ALK and carrying I1171N and a random mutation were seeded into 96-well plates and cultured with gilteritinib (100 nM) for 2 weeks. Ba/F3 cells (20,000 cells) expressing EML4-ALK and carrying G1202R and a random mutation were seeded into 96-well plates and cultured with neladalkib (50 nM) for 2 weeks. Ba/F3 cells (1000 cells) expressing EML4-ALK and carrying I1171N and a random mutation were seeded into 96-well plates and cultured with neladalkib (500 nM) for 2 weeks. Drug-resistant clones were expanded, and the regions encoding the ALK kinase domain were then amplified from the resistant mutants using KOD FX Neo polymerase (TOYOBO, Osaka, Japan). Mutations were detected by Sanger sequencing. For the cell viability assay, Ba/F3 cells expressing EML4–ALK and harboring the predicted resistance mutation were established as described in Section 2.6. Base substitutions were replicated according to the results of Sanger sequencing. For retrovirus infection, the initial culture size was reduced to 2 mL, and instead of spinfection, Ba/F3 cells (4 × 10^5^ cells) were cultured for 24 h in a mixture of 1 mL of RPMI-1640 medium and 1 mL of retrovirus-containing medium supplemented with 8 µg/mL polybrene.

### 2.6 Establishment of Mutated EML4–ALK-expressing Ba/F3 Cells

The predicted resistant mutations were generated in the pMXs-GW-IRES-Puro vector containing the cDNA of EML4–ALK variant 1 plus the G1202R or I1171N mutation using the PrimeSTAR^®^ Mutagenesis Basal Kit (TaKaRa Bio Inc., Shiga, Japan). Subsequently, the product was transformed into competent DH5α *Escherichia coli* cells via 1 min of heat shock at 42℃. The *E. coli* cells were incubated in super optimal broth with catabolite repression medium (2% tryptone, 0.5% yeast extract, 10 mM NaCl, 2.5 mM KCl, 20 mM MgSO4, and 20 mM glucose) for 1 h with intense shaking and then seeded on lysogeny broth (LB) agar plates supplemented with 100 mg/L ampicillin. After overnight incubation, colonies were individually collected and replicated onto other LB agar plates. Colony PCR was performed against the collected colonies. The regions encoding the ALK kinase domain were amplified from the resistant mutants using KAPA Taq Extra DNA polymerase (KAPA Biosystems, Potters Bar, UK), and individual base substitutions were confirmed via Sanger sequencing. Plasmid DNA was prepared using the FastGene™ Plasmid Mini Kit (NIPPON Genetics Co., Ltd, Tokyo, Japan).

### 2.7 Cell Viability Assay

Ba/F3 cells were seeded into 96-well plates (2000 or 4000 cells/well, 90 μL), and 10 μL of serially diluted inhibitors were added to the culture. Three wells were prepared to evaluate cell viability at each drug concentration. Three additional wells were prepared for negative controls by adding 90 μL of the medium and 10 μL of DMSO. After a 72-h incubation period at 37℃, each culture was mixed with 10 μL of Cell Counting Kit-8 (CCK-8) reagent (DOJINDO, Kumamoto, Japan). After 2 h of incubation, the absorbance of the mixture at 450 nm was measured using the Synergy H1 multimode reader (BioTek Instruments, Winooski, VT, USA). The measured score in each well was subtracted from the average score in the negative controls. The score for the 0 nM drug condition was defined as a relative cell viability of 1.0, and the relative cell viability in each well was then calculated. The data were analyzed for drawing graphs, and the half-maximal inhibitory concentration (IC_50_) was then calculated from the data using GraphPad Prism version 6 (GraphPad software, Boston, MA, USA).

### 2.8 Antibodies and Immunoblotting

Ba/F3 cells (1 × 10^6^ cells) were seeded into 12-well plates and treated with different concentrations of inhibitors for 3 h. PBS was used to wash the cells, which were then suspended in TNE buffer (10 mM Tris-HCl [pH 7.4], 1 mM EDTA, 150 mM NaCl, and 1% NP-40). The total protein concentration was measured using the Pierce™ BCA Protein Assay Kit (Thermo Fisher Scientific, Waltham, MA, USA). The suspension was then treated with 2× sample buffer containing 100 mM Tris-HCl (pH 6.8), 4% sodium dodecyl sulfate (SDS), 20% glycerol, 10% 2-mercaptoethanol, and 0.01% bromophenol blue. The samples were thoroughly sonicated and boiled at 95℃ for 5 min. Furthermore, 5 μg of the total protein of the samples were electrophoresed in 7.5% SDS-polyacrylamide gels. The protein was transferred from the gels to Immobilon PVDF membranes (Merck Millipore Ltd., Burlington, MA, USA). The membranes were immersed in appropriate blocking buffer (Tris-buffered saline with 0.05% Tween 20 [TBST] supplemented with 5% w/v bovine serum albumin or 5% w/v nonfat dry milk) for 1 h at room temperature. Subsequently, the membranes were incubated overnight at 4℃ with gentle agitation in primary antibody dilution buffer (phosphorylated ALK [Y1604; Cell Signaling Technologies, Danvers, MA, USA; #3341, 1:1000], ALK [Cell Signaling Technologies; #3791, 1:2000], and α-tubulin [FUJIFILM Wako; #013-25033, 1:4000]). After washing with TBST, the membranes were incubated for 1 h at room temperature with gentle agitation in blocking buffer supplemented with appropriate horseradish peroxidase (HRP)-linked secondary antibodies (anti-rabbit immunoglobulin G [IgG], HRP-linked antibody [Cell Signaling Technologies; #7074, 1:4000], and anti-mouse IgG, HRP-linked antibody [Cell Signaling Technologies; #7076, 1:4000]). The membranes were then washed with TBST and incubated with Immobilon® Western Chemiluminescent HRP Substrate (Merck Millipore Ltd.) or ImmunoStar® LD (FUJIFILM Wako) at room temperature for 3 min. Protein was detected using the ChemiDoc XRS+ System (Bio-Rad Laboratories, Hercules, CA, USA).

### 2.9 Docking Simulation and Molecular Dynamics Simulation

The docking simulation was conducted using the tool sievgene_M included in Medicinally Yielding Protein Engineering SimulaTOr (myPresto) version 5 [20]. The protein coordinate file was prepared by taking the structure from ALK (PDB ID: 4CLI [Structure of the Human Anaplastic Lymphoma Kinase in Complex with PF-06463922: (10R)-7-amino-12-fluoro-2,10,16-trimethyl-15-oxo-10,15,16,17-tetrahydro-2H-8,4-(metheno)pyrazolo(4,3-h)(2,5,11)benzoxadiazacyclotetradecine-3-carbonitrile]). This structure represents the complex of ALK and lorlatinib; therefore, the structure without lorlatinib was used as the template ALK. The chemical structures of the compounds used in this study, namely zotizalkib, gilteritinib, and neladalkib, were manually constructed using Chem3D and subjected to geometry optimization. The protein binding pocket was defined by the makepoint tool, and probe points were generated according to the coordinates of lorlatinib in the original ALK‒lorlatinib complex structure. The docking simulation was then conducted using sievgene_M, and the top 30 docking pose candidates were extracted.

The molecular dynamics simulation was conducted using the tool cosgene included in myPresto version 5 from the acquired docking poses [21]. The gaff21 database was used to assign the molecular force field of the drug to create the topology file. After integrating the protein and drug coordinate files, water molecules and Na and Cl ions were added at a margin of 5 Å from the protein surface, and the tplgeneX tool was used to create topology and coordinate files for the entire system. The Amber parm99 database was used to assign the molecular force field. The boundary condition was determined, and the position of the heavy atoms in the main chain of the protein was constrained for the relief of calculations. After that, the cosgene tool was used to minimize the energy by the steepest descent method to compensate for the strain in the system. The SHAKE method was applied to constrain the interatomic distances of bonds containing hydrogen atoms, and molecular dynamics simulations were performed using the cosgene tool with a time step at 2 fs by 500,000 steps (1 ns in total), and the final structure was obtained.

## 3 Results

### 3.1 Sensitivity of Alectinib-resistant Mutants to Zotizalkib, Gilteritinib, and Neladalkib

Alectinib is firmly established as the standard first-line treatment for ALK-positive NSCLC in Japan. However, disease progression inevitably occurs in many patients because of the emergence of resistance mutations. To evaluate the potential of emerging ALK inhibitors as second-line therapies, we first assessed their efficacy against common alectinib-resistant mutations. We established Ba/F3 cells expressing EML4– ALK variant 1 harboring either the G1202R or I1171N mutation and tested their sensitivity to three next-generation ALK-TKIs: zotizalkib, gilteritinib, and neladalkib (Fig. 1a–c). CCK-8 cell viability assays revealed that the G1202R mutant was sensitive to zotizalkib and neladalkib but resistant to gilteritinib, whereas the I1171N mutant was sensitive to gilteritinib and neladalkib but resistant to zotizalkib (Fig. 1d–f, Table 1), consistent with previous reports [18, 22–24]. Immunoblotting confirmed that zotizalkib and gilteritinib inhibited ALK phosphorylation in cells carrying the G1202R and I1171N mutations, respectively, whereas neladalkib inhibited ALK phosphorylation in both cell lines (Fig. 1g–i). Based on these results, we selected all three drugs for further evaluation as potential sequential treatment options following alectinib.

**Fig. 1.**
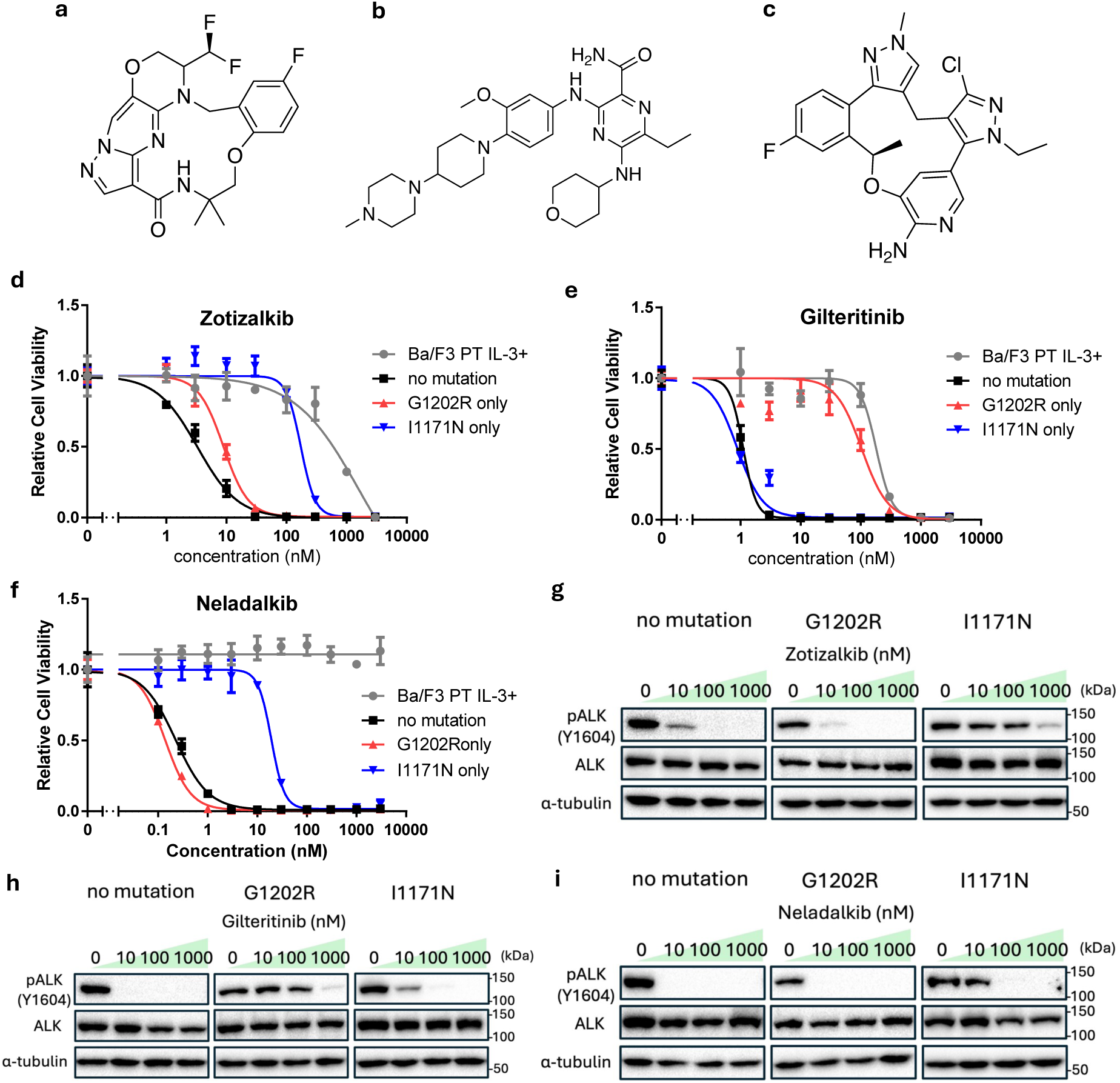
The sensitivity of alectinib-resistant mutants against zotizalkib, gilteritinib, and neladalkib. **a** Chemical structure of zotizalkib. **b** Chemical structure of gilteritinib. **c** Chemical structure of neladalkib. **d–f** Sensitivity evaluation of alectinib-resistant mutants against zotizalkib (**d**), gilteritinib (**e**), and neladalkib (**f**). Ba/F3 cells expressing EML4–ALK variant 1 (v1) plus G1202R or I1171N were exposed to each inhibitor for 72 h. Cell viability was evaluated by CCK-8, with absorbance measured at 450 nm. **g–i** Immunoblotting evaluation of the suppression of phosphorylated ALK expression in alectinib-resistant mutants by zotizalkib (**g**), gilteritinib (**h**), or neladalkib (**i**). Ba/F3 cells expressing EML4–ALK v1 and each resistance mutation were treated with different inhibitors for 3 h. Next, immunoblotting was used to detect the indicated protein in cell lysates. *CCK-8* Cell Counting Kit-8, *EML4* echinoderm microtubule-associated protein-like 4, *ALK* anaplastic lymphoma kinase

**Table 1.** IC_50_s of ALK-TKIs in Ba/F3 cells carrying alectinib-resistant mutations.

| IC<sub>50</sub> (nM) |  |  |  |
| --- | --- | --- | --- |
|  | Zotizalkib | Gilteritinib | Neladalkib |
| Ba/F3 PT IL-3+ | 1630 | 182.6 | N.D. |
| no mutation | 6.5 | 1.1 | 0.2 |
| G1202R | 9.0 | 107.1 | 0.1 |
| I1171N | 190.7 | 0.9 | 19.1 |

### 3.2 Experimental System for Predicting Resistance Mutations Using Error-prone PCR

Acquired point mutations in the ALK kinase domain represent the primary mechanism of resistance following ALK-TKI therapy. Therefore, predicting resistance mutations at the preclinical stage is important for developing effective therapies following alectinib treatment. To address this, we previously established a novel preclinical prediction platform using error-prone PCR [18]. In the present study, the cDNA libraries were transfected into Plat-E cells, packaged into retroviruses, and used to infect 6 × 10^7^ Ba/F3 cells. After puromycin selection, the estimated infection efficiencies were 3.44% (zotizalkib, G1202R), 2.53% (gilteritinib, I1171N), 21.4% (neladalkib, G1202R), and 10.8% (neladalkib, I1171N), as presented in Supplementary Fig. 1a. As the cDNA mutant library was constructed from approximately 7 × 10^5^ colonies and that more than 1 × 10^6^ Ba/F3 cells were infected per condition, it is likely that the full spectrum of possible point mutations was represented in the transduced cell populations.

### 3.3 Prediction of Zotizalkib-Resistant Mutations and Sensitivity to Other ALK-TKIs

First, we predicted resistance mutations to zotizalkib. A Ba/F3 mutant library, in which each cell harbored the G1202R mutation along with a random secondary mutation, was treated with 100 nM zotizalkib for 2 weeks. Under these conditions, sensitive cells were expected to be completely eliminated, as indicated by the results of the cell viability assay for alectinib-resistant mutants treated with zotizalkib (Fig. 1d). Cells were seeded at a low density into 96-well plates to isolate individual resistant clones. After 2 weeks of drug exposure, Sanger sequencing of the surviving clones revealed several compound mutations associated with zotizalkib resistance, including G1202R + F1174L/C/I/V, G1202R + F1245V, and G1202R + P1153H (Fig. 2a). Clinically, the compound mutation G1202R + F1174L has been reported as a lorlatinib-resistant mutation [25]. A point mutation at F1245 has been identified in patients with neuroblastoma [26]. Meanwhile, the P1153H mutation has not been previously reported.

**Fig. 2.**
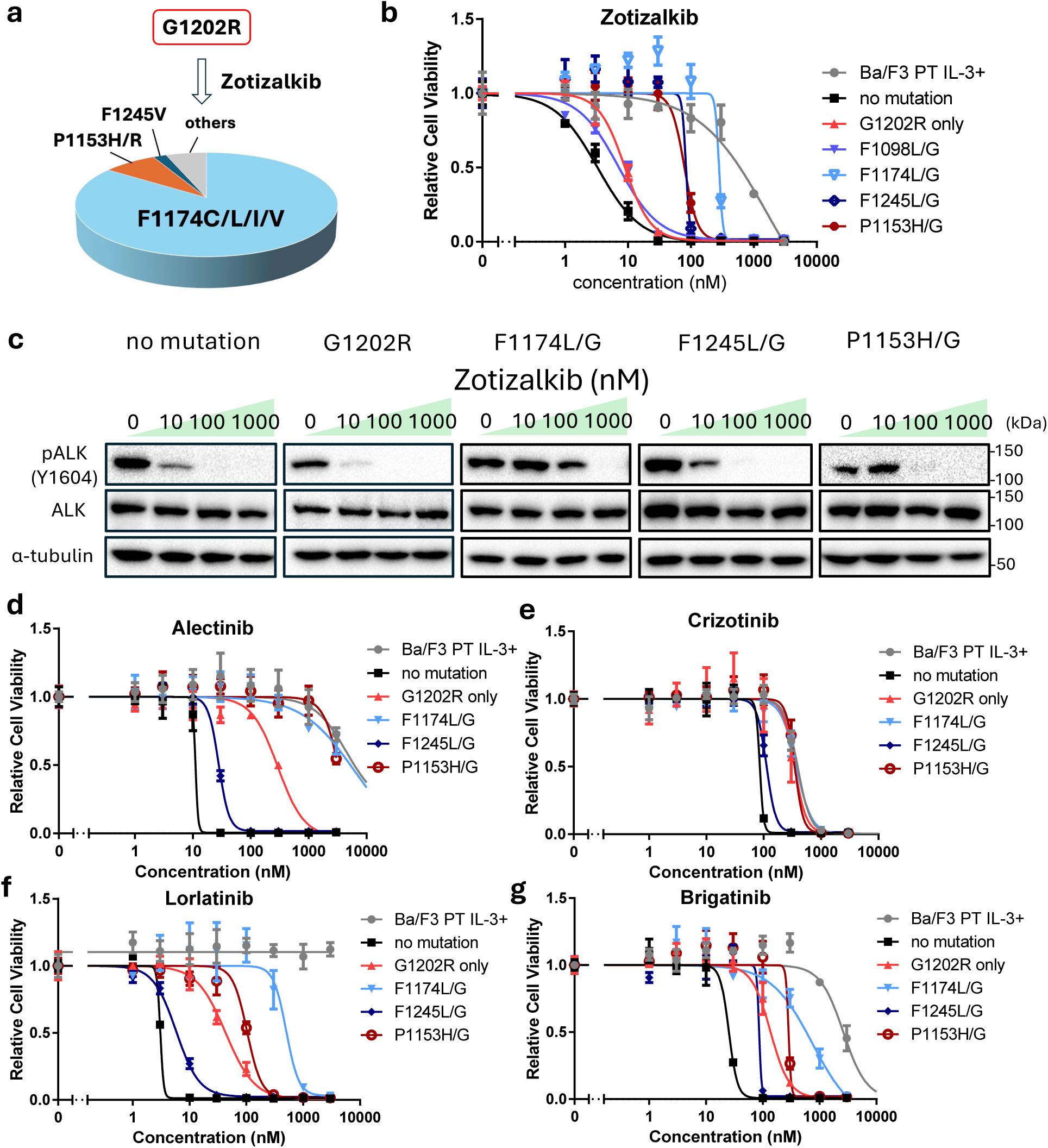
Zotizalkib-resistant mutations emerging from G1202R-positive ALK and their cross-sensitivity to other ALK-TKIs. **a** Predicted zotizalkib-resistant mutations (n = 54). Mutant libraries of Ba/F3 cells were exposed to zotizalkib (100 nM) for 2 weeks. **b** Sensitivity evaluation of predicted zotizalkib-resistant mutants against zotizalkib. Ba/F3 cells expressing EML4–ALK variant 1 and a zotizalkib-resistant mutation were exposed to zotizalkib for 72 h. Cell viability was evaluated using the CCK-8 assay, with absorbance measured at 450 nm. **c** Immunoblotting evaluation of the suppression of phosphorylated ALK expression in predicted zotizalkib-resistant mutants by zotizalkib. Ba/F3 cells expressing EML4–ALK variant 1 and a resistance mutation were treated with zotizalkib for 3 h. Next, immunoblotting was used to detect the indicated protein in cell lysates. **d–g** Sensitivity evaluation of predicted zotizalkib-resistant mutants against alectinib (**d**), crizotinib (**e**), lorlatinib (**f**), and brigatinib (**g**). Ba/F3 cells expressing EML4–ALK variant 1 with each resistance mutation were exposed to zotizalkib for 72 h. Cell viability was evaluated by CCK-8 and absorbance at a wavelength of 450 nm. *CCK-8* Cell Counting Kit-8, *EML4* echinoderm microtubule-associated protein-like 4, *ALK* anaplastic lymphoma kinase, *TKIs* tyrosine kinase inhibitors

Next, we investigated whether the predicted point mutations conferred resistance to zotizalkib. Ba/F3 cells expressing EML4–ALK and each of the predicted zotizalkib-resistant mutations were established, and their drug sensitivity was evaluated. These additional point mutations conferred greater resistance to zotizalkib than G1202R alone (Fig. 2b, c). To identify potential therapeutic strategies following zotizalkib treatment, we further evaluated the sensitivity of the acquired zotizalkib-resistant mutants to various ALK-TKIs. Both the G1202R + F1174L and G1202R + P1153H mutants were resistant to alectinib (Fig. 2d), crizotinib (Fig. 2e), lorlatinib (Fig. 2f), and brigatinib (Fig. 2g), whereas the G1202R + F1245L mutant remained sensitive to all of these ALK-TKIs.

### 3.4 Prediction of Gilteritinib-resistant Mutations and Sensitivity to Other ALK-TKIs

Next, we predicted gilteritinib-resistant mutations using the same procedure described in Section 3.3. A Ba/F3 mutant library, in which each cell harbored the I1171N mutation along with a random secondary mutation, was treated with 100 nM gilteritinib for 2 weeks. The compound mutations I1171N + E1210K/A, I1171N + E1129V, and I1171N + D1203N were identified as gilteritinib-resistant mutations (Fig. 3a). Single-point mutations at E1129, E1210, and D1203 were previously reported in patients who developed resistance to ALK-TKIs [27, 28]. Using the same validation approach described in Section 3.3, we confirmed that these additional point mutations increased resistance to gilteritinib compared with the effects of I1171N alone (Fig. 3b, c). Furthermore, these resistant mutants exhibited cross-resistance to other ALK-TKIs (Fig. 3d–g).

**Fig. 3.**
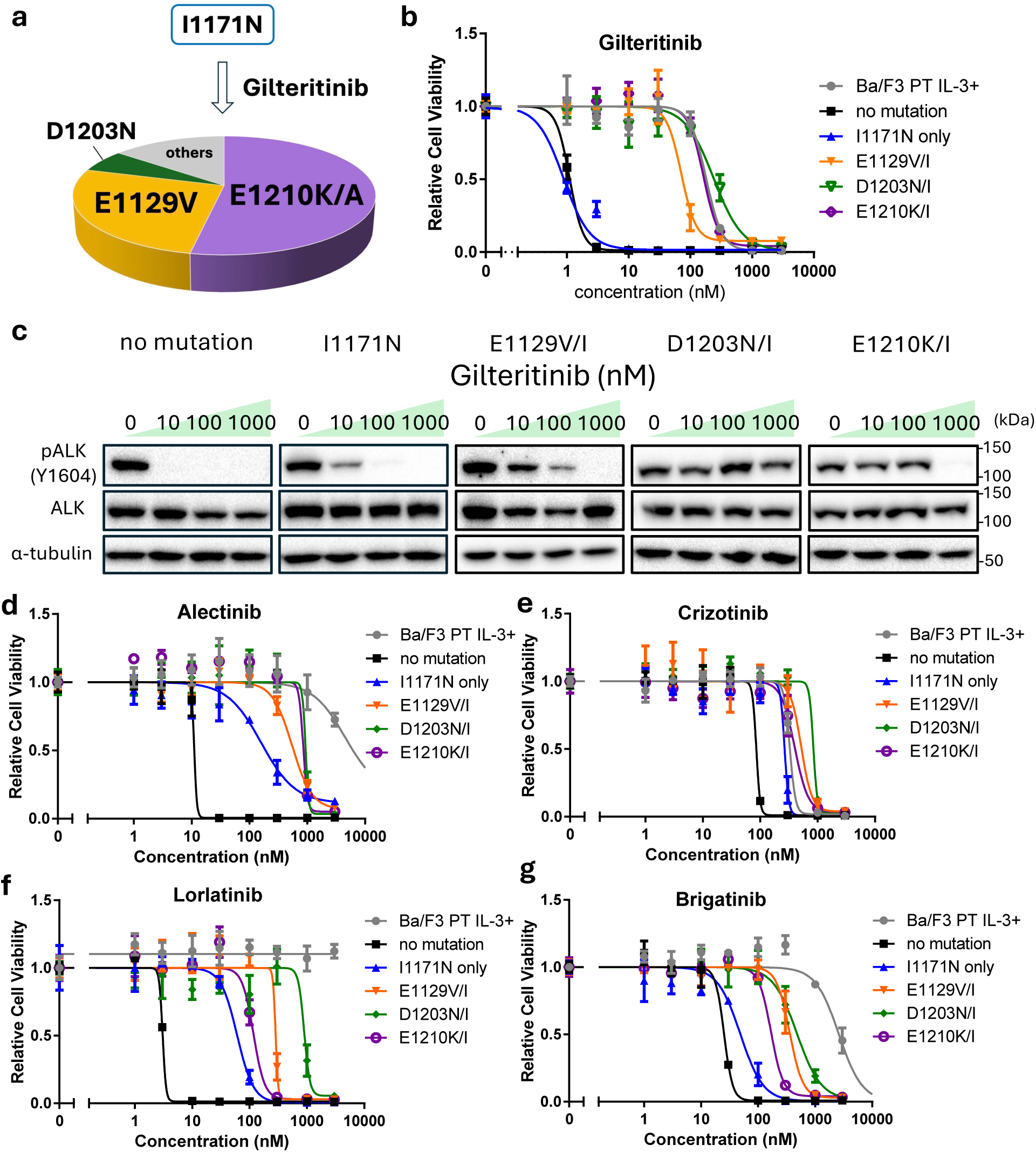
Gilteritinib-resistant mutations emerging from I1171N-positive ALK and their cross-sensitivity to other ALK-TKIs. **a** Predicted gilteritinib-resistant mutations (n = 49). Mutant libraries of Ba/F3 cells were exposed to gilteritinib (100 nM) for 2 weeks. **b** Sensitivity of the predicted resistant mutants to gilteritinib. Ba/F3 cells expressing EML4–ALK variant 1 and each mutation were exposed to gilteritinib for 72 h. Cell viability was evaluated by the CCK-8 assay, with absorbance measured at 450 nm. **c** Immunoblotting evaluation of the suppression of phosphorylated ALK expression in each resistant mutant by gilteritinib. Gilteritinib was given to Ba/F3 cells expressing EML4-ALK variant 1 with each resistance mutation for 3 h. Next, immunoblotting was used to detect the indicated protein in cell lysates. **d–g** Sensitivity evaluation of each resistance mutants against alectinib (**d**), crizotinib (**e**), lorlatinib (**f**), and brigatinib (**g**). Ba/F3 cells expressing EML4-ALK variant 1 with each resistance mutation were exposed to zotizalkib for 72 h. Cell viability was evaluated by the CCK-8 assay, with absorbance measured at 450 nm. *CCK-8* Cell Counting Kit-8, *EML4* echinoderm microtubule-associated protein-like 4, *ALK* anaplastic lymphoma kinase, *TKIs* tyrosine kinase inhibitors

### 3.5 Prediction of Neladalkib-resistant Mutations and Sensitivity to Other ALK-TKIs

Finally, we predicted resistance mutations to neladalkib using the procedure described in Sections 3.3 and 3.4. We first screened a mutant Ba/F3 cell library harboring the G1202R mutation along with a random secondary mutation. However, even at reduced drug concentrations, extremely few cells survived. At 50 nM, only seven resistant clones were isolated, all of which harbored the G1202R + L1196M compound mutation (Fig. 4a, b). Based on the sensitivity assays, the G1202R + L1196M mutant remained responsive to neladalkib, suggesting that no additional mutations combined with G1202R conferred resistance to this compound.

**Fig. 4.**
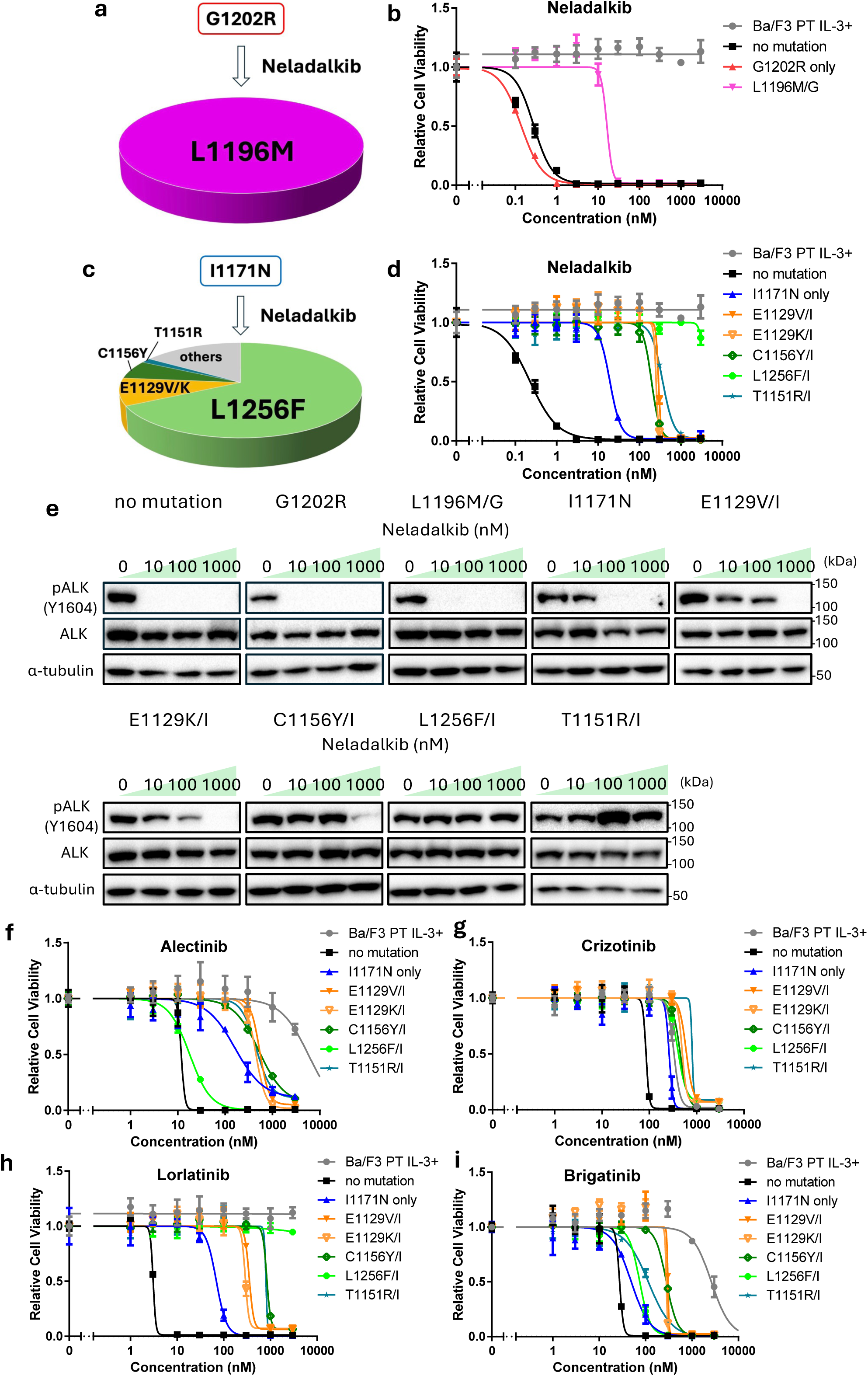
Neladalkib-resistant mutations emerging from G1202R- or I1171N-positive ALK and their cross-sensitivity to other ALK-TKIs. **a** Predicted neladalkib-resistant mutations with G1202R (n = 7). Mutant libraries of Ba/F3 cells were exposed to neladalkib (50 nM) for 2 weeks. **b** Sensitivity evaluation of the G1202R + L1196M compound variant against neladalkib. Ba/F3 cells expressing EML4–ALK variant 1 and different mutants were exposed to neladalkib for 72 h. Cell viability was evaluated by the CCK-8, with absorbance measured at 450 nm. **c** Predicted neladalkib-resistant mutations with I1171N (n = 64). Mutant libraries of Ba/F3 cells were exposed to neladalkib (500 nM) for 2 weeks. Other mutations included the triple mutation I1171N + C1156Y + D1203N, as well as I1171N + L1196M and I1171N + V1130L, both of which remained sensitive to neladalkib. **d** Sensitivity of predicted mutations to neladalkib. Ba/F3 cells expressing EML4– ALK variant 1 and different mutants were exposed to neladalkib for 72 h. Cell viability was evaluated by the CCK-8 assay, with absorbance measured at 450 nm. **e** Immunoblotting evaluation of the suppression of phosphorylated ALK expression in each resistant mutant by neladalkib. Ba/F3 cells expressing EML4– ALK variant 1 and different resistance mutations were treated with neladalkib for 3 h. Next, immunoblotting was used to detect the indicated protein in cell lysates. **f–i** Sensitivity of each resistance mutant to alectinib (**f**), crizotinib (**g**), lorlatinib (**h**), and brigatinib (**i**). Ba/F3 cells expressing EML4–ALK variant 1 and different resistance mutations were exposed to neladalkib for 72 h. Cell viability was evaluated by the CCK-8 assay, with absorbance measured at 450 nm. *CCK-8* Cell Counting Kit-8, *EML4* echinoderm microtubule-associated protein-like 4, *ALK* anaplastic lymphoma kinase, *TKIs* tyrosine kinase inhibitors

Conversely, screening a Ba/F3 mutant library harboring I1171N and a random mutation at a drug concentration of 500 nM revealed several candidate resistance mutations, including I1171N + L1256F, I1171N + E1129V/K, I1171N + C1156Y, and I1171N + T1151R (Fig. 4c). The I1171N + L1256F mutation was reported as a lorlatinib-resistant mutation in an ENU-based mutagenesis screen, whereas the I1171N + C1156Y and I1171N + T1151R mutations were identified in clinical samples [10, 29, 30].

To evaluate whether these predicted point mutations conferred resistance to neladalkib, we established Ba/F3 cells expressing EML4–ALK with each mutation and assessed their drug sensitivity. These additional point mutations increased resistance to neladalkib compared with the effects of the I1171N alone (Fig. 4d, e). Furthermore, these resistant mutants exhibited cross-resistance to all tested ALK-TKIs (Fig. 4f–i).

## 4 Discussion

### 4.1 Mechanisms of Drug Resistance Associated with Each Mutation

The ALK kinase domain adopts a typical bilobed kinase fold consisting of an N-lobe, C-lobe, and intervening hinge region. The N-lobe contains a five-stranded β-sheet, the αC-helix involved in activation, and a phosphate-binding loop (P-loop) that interacts with ATP. By contrast, the C-lobe harbors the activation loop (A-loop) and catalytic loop, both of which are critical for catalytic activity. ATP binds via hydrogen bonding between its adenine ring and the hinge region, as well as through interactions with the P-loop, αC-helix, and DFG motif within the A-loop [31–34] (Fig. 5a). We subsequently examined how each resistance-associated mutation altered these structural elements and thereby reduces the efficacy of individual ALK inhibitors.

**Fig. 5.**
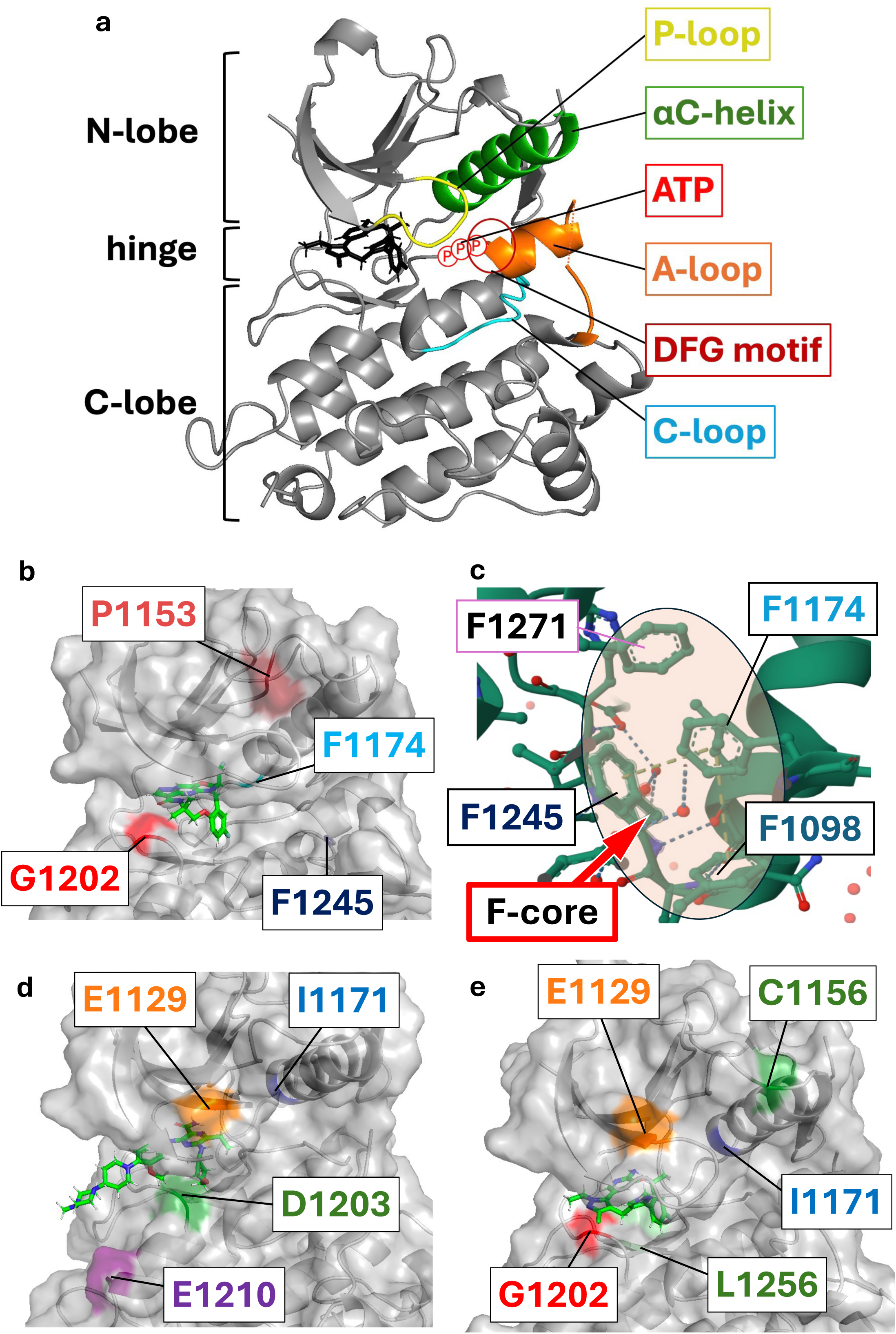
Structural mapping of resistance mutations on the ALK kinase domain **a** Structure of the ALK kinase domain highlighting key activation-related regions. **b** Docking pose of zotizalkib in the ALK kinase domain. **c** Spatial configuration of the phenylalanine core within the ALK kinase domain. **d** Docking pose of gilteritinib in the ALK kinase domain. **e** Docking pose of neladalkib in the ALK kinase domain. *ALK* anaplastic lymphoma kinase

#### 4.1.1 Zotizalkib-resistant Mutations

Zotizalkib exhibited potent activity against the G1202R mutation, but it was ineffective against I1171N (Fig. 1d, g). The G1202R mutation introduces a bulky arginine residue that protrudes from the hinge region into the inhibitor-binding site, resulting in steric hindrance and resistance to inhibitors such as alectinib [35]. However, zotizalkib contains a similar macrocyclic structure as lorlatinib and binds without extending into the hinge region, likely accounting for its retained efficacy against the G1202R mutant. Conversely, I1171 is located deep within the ATP-binding pocket on the αC-helix. Mutations at this site have been reported to induce positional shifts in the adjacent residue E1167, thereby distorting the αC-helix and permitting inhibitors to weaken hydrogen bonds, thereby conferring resistance [35]. Because zotizalkib probably interacts deeply within the pocket through hydrogen bonding and other interactions, the I1171N mutation likely disrupts these interactions, resulting in a loss of binding and diminished efficacy.

In this study, G1202R + F1174C/L/I/V, G1202R + F1245L/V, and G1202R + P1153H were identified to confer resistance to zotizalkib (Fig. 5b). In the inactive conformation of ALK, the DFG motif residue F1271, together with the neighboring phenylalanines F1098, F1174, and F1245, forms a hydrophobic aromatic cluster called the F-core, which stabilizes the kinase in its inactive state (Fig. 5c) [36]. Thus, mutations at F1174 and F1245, which remove the aromatic side chains, might destabilize the F-core and promote a shift toward the active conformation. To test this hypothesis, we introduced leucine substitutions at these phenylalanine residues (excluding F1271, as mutation of this residue would likely generate a kinase-dead protein because of it is located within the catalytically critical DFG motif [37]). Drug sensitivity assays revealed that only the F1174L and F1245L mutants, but not the F1098L mutant, conferred resistance to zotizalkib (Fig. 2b). These results suggest that F1174, F1245, and F1271 are essential for maintaining the F-core, whereas F1098, located more distally from F1271, has a less critical role. Finally, although previously unreported, the P1153H mutation is positioned between the β-sheet of the N-lobe and the αC-helix, suggesting that it might distort one or both structural elements.

#### 4.1.2 Gilteritinib-resistant Mutations

Gilteritinib was effective against I1171N but not against G1202R (Fig. 1e, h). Its shallow binding mode likely enables it to retain activity against I1171N. However, because gilteritinib also binds by covering the hinge region, the steric hindrance introduced by the bulky arginine residue at residue 1202 likely impairs binding and renders the drug ineffective.

In this study, we identified I1171N + E1210K, I1171N + E1129V, and I1171N + D1203N as compound mutations that confer resistance to gilteritinib (Fig. 5d). The E1210K mutation replaces a negatively charged glutamic acid with a positively charged lysine, likely converting electrostatic attraction into repulsion and thereby inhibiting drug binding. E1129 is located near the P-loop, and it forms hydrogen bonds with surrounding residues such as H1124 and T1151, stabilizing the P-loop [38]. The E1129V mutation likely disrupts these interactions, thereby destabilizing the P-loop and conferring resistance to gilteritinib. D1203 forms a stabilizing hydrogen bond with a nearby serine residue (S1206) and helps maintain the α-helical structure [39]. A change in the charge at this site could disrupt this interaction and destabilize the helix, ultimately leading to drug resistance.

#### 4.1.3 Neladalkib-resistant Mutations

Neladalkib was effective against both the G1202R and I1171N mutations (Fig. 1f, i). Similarly as zotizalkib, it possesses a macrocyclic structure and binds without extending into the hinge region, which likely explains its efficacy against the G1202R mutant. Compared with the three tandem macrocycles of zotizalkib, neladalkib’s macrocyclic–carbon–macrocyclic configuration might offer greater flexibility, allowing it to accommodate conformational distortions in deeper regions of the ALK kinase domain, such as I1171N.

In this study, we identified I1171N + L1256F, I1171N + C1156Y, I1171N + E1129V/K, and I1171N + T1151R as compound mutations conferring resistance to neladalkib (Fig. 5e). The I1171N + L1256F mutation, previously reported to confer resistance to lorlatinib [30], also induced resistance to neladalkib, which shares a similar binding mode. This resistance is likely attributable to steric hindrance between the phenyl ring of F1256 and the six-membered ring of neladalkib, as previously reported for L1256F-mediated resistance to lorlatinib [40]. C1156 is located at the N-terminal end of the αC-helix. The C1156Y mutation introduces a bulky tyrosine residue that protrudes toward the P-loop, likely disrupting the stability of the P-loop and leading to increased kinase activity and drug resistance [41]. On the contrary, gilteritinib retained efficacy against these mutations [18], possibly because of its multiple flexible rings and the presence of additional amino and amide groups capable of forming hydrogen bonds with the hinge region. For compound mutations such as I1171N + E1129V/K and I1171N + T1151R, resistance might similarly arise from the loss of hydrogen bonding among E1129, H1124, and T1151, resulting in P-loop destabilization, as observed with gilteritinib [38].

### 4.2 Sequential ALK-TKI Strategies for Overcoming Resistance in Compound ALK Mutations Following Alectinib Failure

The clinical introduction of multiple ALK-TKIs, including alectinib and lorlatinib, has markedly improved outcomes for patients with ALK-positive NSCLC [9]. However, disease relapse attributable to resistance mutations remains a major challenge, particularly after lorlatinib treatment [13]. As resistance profiles vary depending on the ALK-TKI used, mutation-guided sequential treatment strategies are critically needed to sustain therapeutic efficacy.

To address this issue, predicting resistance mutations that emerge after specific ALK-TKI treatments and identifying drugs effective against these variants represent a promising approach. In this study, we applied our previously established resistance mutation prediction method based on error-prone PCR [18]. Using this method, we examined the novel ALK inhibitors zotizalkib, gilteritinib, and neladalkib as second-line therapies following alectinib resistance. Zotizalkib was effective against G1202R, gilteritinib was effective against I1171N, and neladalkib was effective against both mutations (Fig. 1). Based on these results, we predicted potential compound mutations that might arise during hypothetical second-line treatment with each drug (Figs. 2–4).

We then evaluated the sensitivity of these predicted resistance variants to each inhibitor. Although most of the variants exhibited resistance to currently approved ALK-TKIs, all variants were effectively targeted by at least one of the three investigational drugs. For compound mutations involving G1202R, neladalkib consistently exhibited potent activity, suggesting that treatment with neladalkib after G1202R arises following alectinib failure could help sustain therapeutic efficacy.

For compound mutations involving I1171N, there was no overlap in resistance profiles between gilteritinib and neladalkib, implying that sequential or combinational administration of these agents could maintain efficacy. Notably, although the I1171N + E1129V variant was predicted to confer gilteritinib resistance, its IC_50_ was 68.9 nM (Table 2). Conversely, G1202R or I1171N mutants treated with lorlatinib had similar IC_50_s (approximately 90 nM), for which clinical responses have been reported [14]. Thus, this variant might have remained moderately sensitive to gilteritinib. In our previous study, we identified E1129V, C1156Y, and L1256F as secondary mutations that confer resistance to ensartinib when acquired in addition to I1171N [18]. These mutations were also associated with resistance to neladalkib; however, gilteritinib remained effective against all of these variants. Furthermore, compound mutations such as I1171N + E1210K and I1171N + D1203N, which confer resistance to gilteritinib, were not identified as resistant clones in our previous screening [18]. These findings suggest that these variants could retain sensitivity to ensartinib, hinting that sequential or combinational administration of ensartinib and gilteritinib could represent an effective treatment strategy for patients harboring I1171N-based resistance mutations.

**Table 2.**
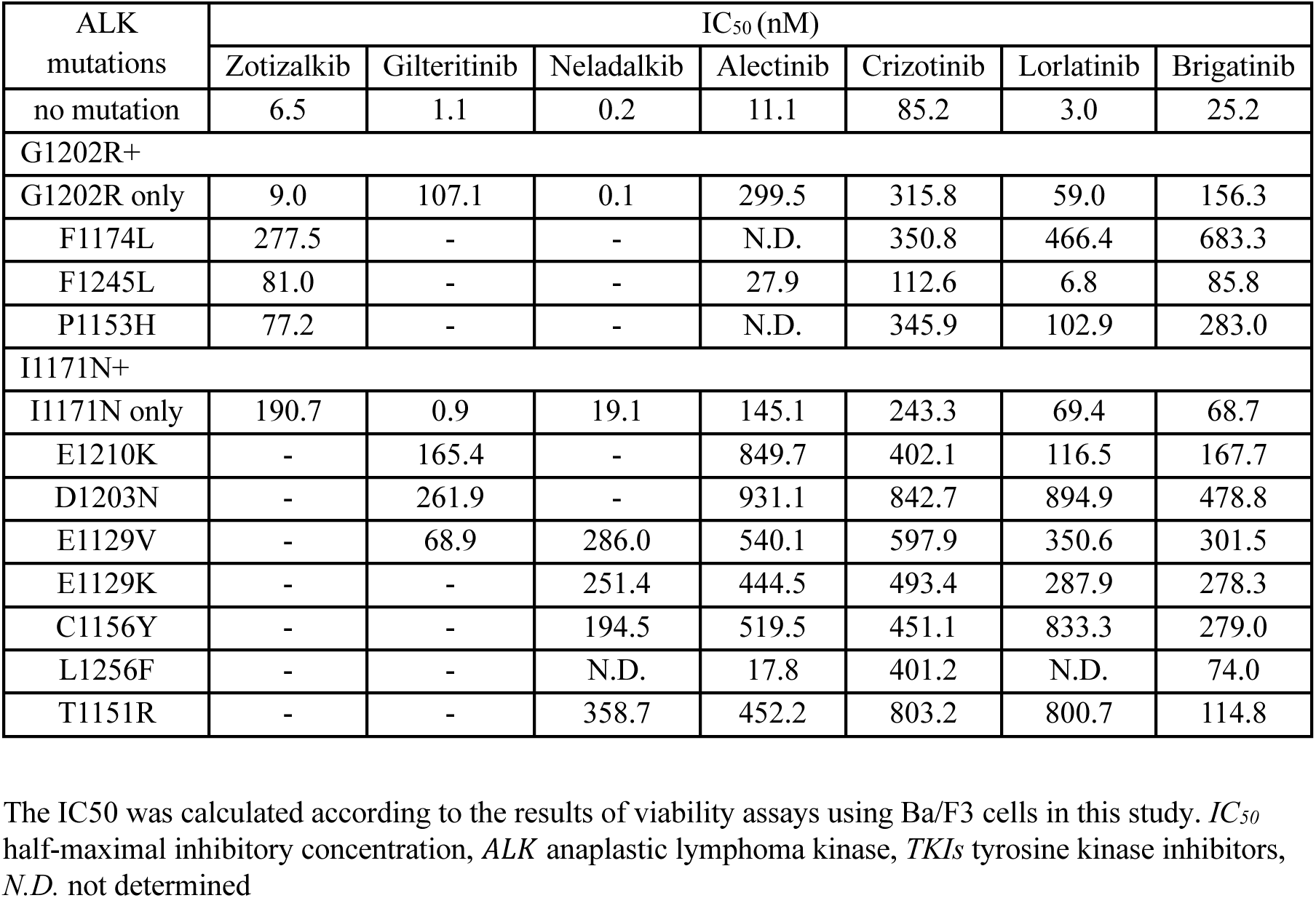
IC_50_s of several ALK-TKIs in the presence of various resistance mutations.

| ALK mutations | IC<sub>50</sub> (nM) |  |  |  |  |  |  |
| --- | --- | --- | --- | --- | --- | --- | --- |
|  | Zotizalkib | Gilteritinib | Neladalkib | Alectinib | Crizotinib | Lorlatinib | Brigatinib |
| no mutation | 6.5 | 1.1 | 0.2 | 11.1 | 85.2 | 3.0 | 25.2 |
| G1202R+ |  |  |  |  |  |  |  |
| G1202R only | 9.0 | 107.1 | 0.1 | 299.5 | 315.8 | 59.0 | 156.3 |
| F1174L | 277.5 | - | - | N.D. | 350.8 | 466.4 | 683.3 |
| F1245L | 81.0 | - | - | 27.9 | 112.6 | 6.8 | 85.8 |
| P1153H | 77.2 | - | - | N.D. | 345.9 | 102.9 | 283.0 |
| I1171N+ |  |  |  |  |  |  |  |
| I1171N only | 190.7 | 0.9 | 19.1 | 145.1 | 243.3 | 69.4 | 68.7 |
| E1210K | - | 165.4 | - | 849.7 | 402.1 | 116.5 | 167.7 |
| D1203N | - | 261.9 | - | 931.1 | 842.7 | 894.9 | 478.8 |
| E1129V | - | 68.9 | 286.0 | 540.1 | 597.9 | 350.6 | 301.5 |
| E1129K | - | - | 251.4 | 444.5 | 493.4 | 287.9 | 278.3 |
| C1156Y | - | - | 194.5 | 519.5 | 451.1 | 833.3 | 279.0 |
| L1256F | - | - | N.D. | 17.8 | 401.2 | N.D. | 74.0 |
| T1151R | - | - | 358.7 | 452.2 | 803.2 | 800.7 | 114.8 |

The IC<sub>50</sub> was calculated according to the results of viability assays using Ba/F3 cells in this study. *IC<sub>50</sub>* half-maximal inhibitory concentration, *ALK* anaplastic lymphoma kinase, *TKIs* tyrosine kinase inhibitors, *N.D.* not determined

Taken together, our findings suggest that the use of neladalkib for G1202R-positive relapses and sequential use of gilteritinib–neladalkib or gilteritinib–ensartinib for I1171N-positive relapses could sustain treatment efficacy and delay the onset of resistance (Fig. 6).

**Fig. 6.**
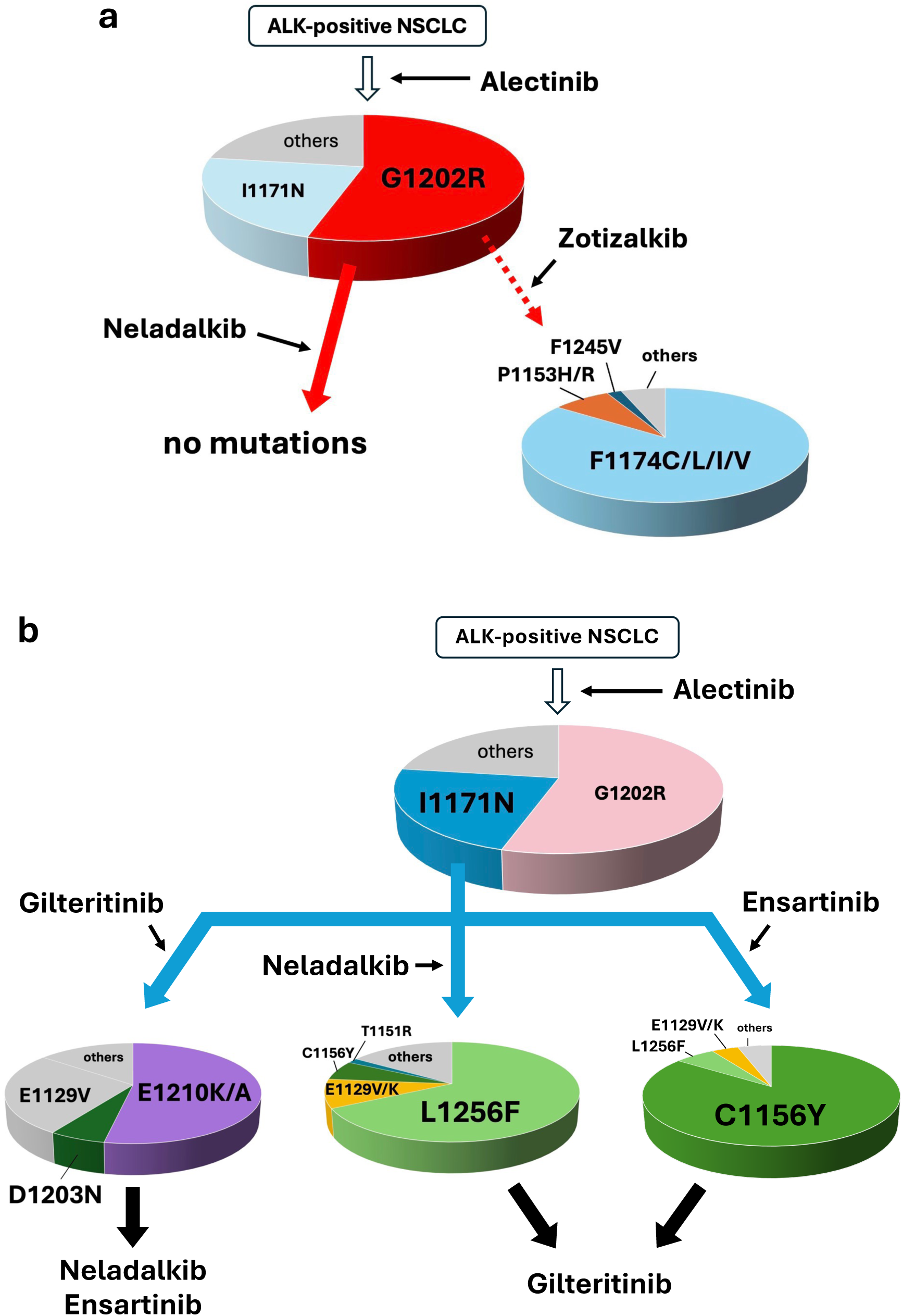
Possible therapeutic strategies after alectinib failure. **a** Neladalkib is available for G1202R-harboring ALK-positive NSCLC, and no mutations were predicted to confer resistance to neladalkib. Zotizalkib is also available for G1202R-harboring ALK-positive NSCLC; however, additional resistance mutations such as F1174C/L/I/V may emerge. Each slice size in the compound mutation pie chart reflects the proportion of mutant clones obtained. **b** Gilteritinib is a treatment option for patients with I1171N-positive ALK-rearranged NSCLC. Most relapses may occur due to an additional E1210K or D1203N mutation. In such cases, neladalkib or ensartinib may be effective. On the other hand, treatment of I1171N-mutant cells with neladalkib or ensartinib may lead to relapse due to the emergence of additional mutations such as L1256F, E1129V, or C1156Y; however, these compound mutants are considered to remain sensitive to gilteritinib. Sequential administration of gilteritinib-neladalkib or gilteritinib-ensartinib is expected to be beneficial for I1171N-positive relapses. Each slice size in the compound mutation pie chart reflects the proportion of mutant clones obtained. *ALK* anaplastic lymphoma kinase, *NSCLC* non-small cell lung cancer

## 5. Conclusion

Using our error-prone PCR-based system, we identified potential mutations conferring resistance to zotizalkib, gilteritinib, or neladalkib. Based on these findings, we propose that these investigational drugs could represent alternative second-line treatment options to lorlatinib following alectinib failure. In clinical settings, resistance mutations are often heterogeneous, and they might not be fully captured by current diagnostic tests. Consequently, physicians might need to empirically select subsequent therapies without definitive molecular guidance. Even when molecular testing is not feasible, selecting drugs with predicted activity against plausible resistance mutations provides a rational basis for treatment decisions. When such resistance mutations can be anticipated, incorporating this information could further enhance the precision and clinical efficacy of sequential ALK-TKI strategies. With the continued development of ALK-TKIs, we believe that individualized clinical strategies—guided by predicted resistance mutation profiles when available—could help optimize therapeutic outcomes while preserving patients’ quality of life.

## Figure Legends

**Sup Fig. 1**

Calculation of infection efficiency **a** Cell growth after puromycin addition to calculate infection efficacy (zotizalkib, gilteritinib). The cell concentration at 0 h was 2 × 10^5^ cells/mL. Living cells were counted at 42, 66, and 93 h after puromycin addition, and regression lines were drawn. The confidence of determination was 0.976 for zotizalkib (G1202R) and 0.997 for gilteritinib (I1171N), and the predicted initial infected cell concentration was 0.688 × 10^5^ cells/mL for zotizalkib (G1202R) and 0.505 × 10^5^ cells/mL for gilteritinib (I1171N). Consequently, the infection efficiency was calculated as 3.44% for zotizalkib (G1202R) and 2.53% for gilteritinib (I1171N). **b** Cell growth observation after puromycin addition to calculate infection efficacy (neladalkib). The cell concentration at 0 h was 2 × 10^5^ cells/mL. Living cells were counted at 54, 68, and 80 h after puromycin addition, and regression lines were drawn. The confidence of determination was 0.937 for G1202R and 0.957 for I1171N, and the predicted initial infected cell concentration was 4.28 × 10^5^ cells/mL for G1202R and 2.16 × 10^5^ cells/mL for I1171N. Consequently, the infection efficiency was calculated as 21.4% for G1202R and 10.8% for I1171N.

**Sup Fig. 2**

The F1245L secondary mutation restores sensitivity to ALK-TKIs in G1202R-positive ALK **a–d** Immunoblotting evaluation of the suppression of phosphorylated ALK expression in the presence of no mutation, G1202R alone, and G1202R + F1245L by alectinib (**a**), crizotinib (**b**), lorlatinib (**c**), or brigatinib (**d**). Ba/F3 cells expressing EML4–ALK variant 1 and different resistance mutation were treated with each inhibitor for 3 h. Next, immunoblotting was used to detect the indicated protein in cell lysates. *EML4* echinoderm microtubule-associated protein-like 4, *ALK* anaplastic lymphoma kinase

## Declarations

### Availability of data and material

The data and material in this study are available from the corresponding author upon reasonable request.

### Ethics approval

Not applicable.

### Consent to participate

Not applicable.

### Consent for publication

Not applicable.

### Code availability

Not applicable.

### Author contributions

Y.D. and K.S. conceived and designed this study. Y.T., H.K., C.A. and Y.D performed the experiments and analyzed the data. Y.T., Y.D and K.S. interpreted the data and wrote the manuscript. All the authors reviewed and edited the manuscript.

## Acknowledgment

The authors would like to thank Enago (www.enago.jp) for the English language review. We appreciate our lab members for their valuable discussions.

