## Supplementary figures and images for "Preclinical Prediction of Resistance and Optimization of Sequential Therapy for ALK-positive Lung Cancer Using Next-generation ALK Inhibitors"

### Supplemental Figures

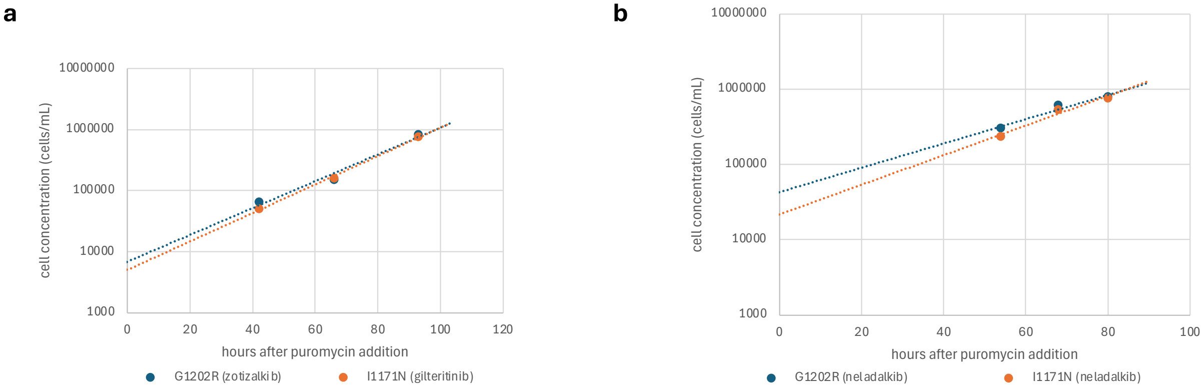

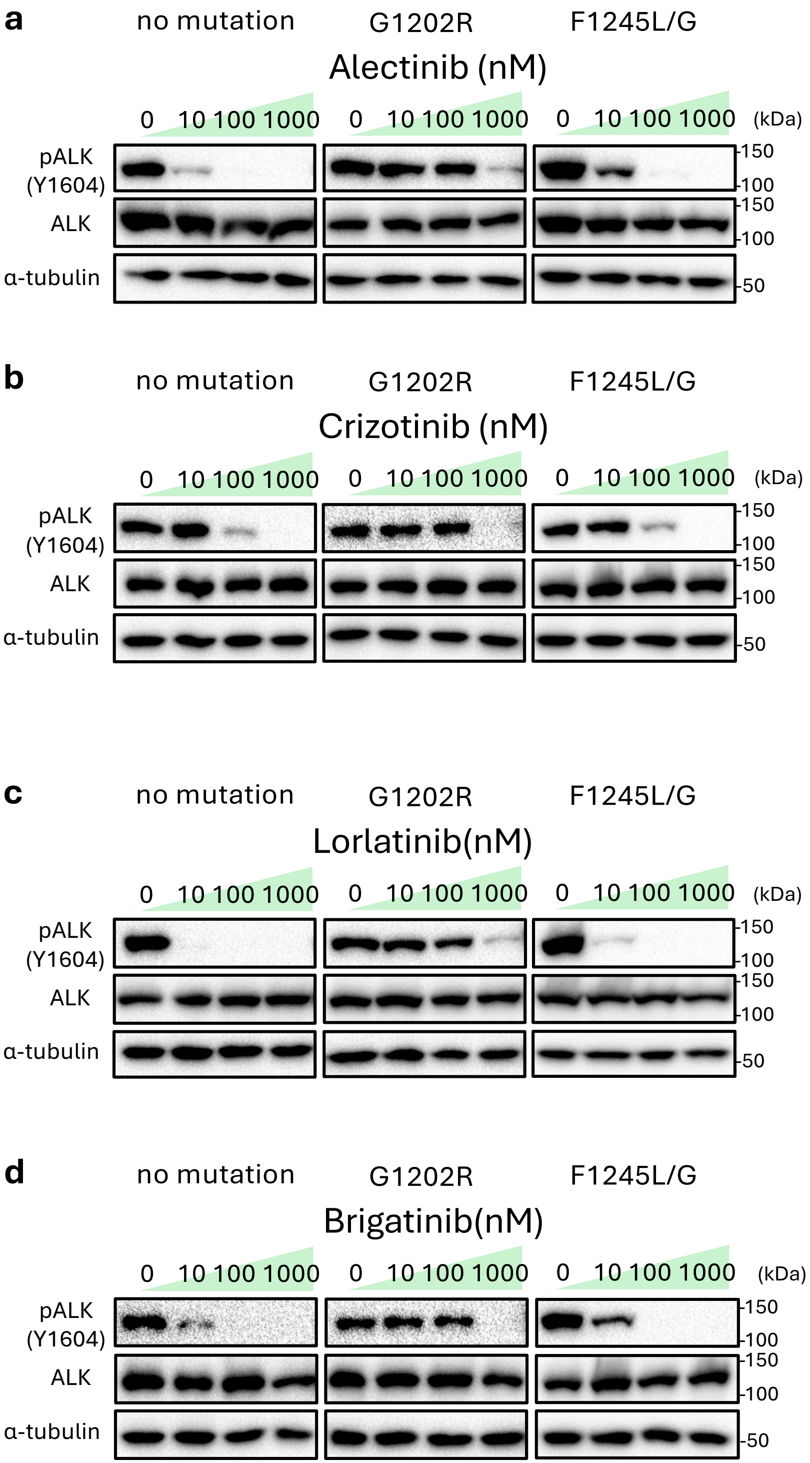
